# Supramodal and modality-specific neural information supports multi-feature prediction errors across cortical levels

**DOI:** 10.1101/2024.02.12.578256

**Authors:** Maria Niedernhuber, Francesca Fardo, Micah Allen, Tristan Bekinschtein

**Author notes:** Correspondence concerning this article should be addressed to Maria Niedernhuber, Department of Psychology, Downing Pl, Cambridge CB2 3EB, United Kingdom.

## Abstract

Predictive coding posits that the brain actively anticipates inputs from different senses, generating prediction errors when incoming information deviates from internal expectations. While much research has focused on prediction errors elicited by violations of single sensory features, natural environments frequently present more complex events deviating across multiple stimulus dimensions and sensory modalities. In this study, we employed a hierarchical oddball paradigm (n=30) manipulating auditory and somatosensory stimuli to violate one or two sensory features while high-density EEG was recorded. Temporal decoding revealed that while both single- and double-deviants evoked sustained supramodal activation patterns, double-deviants uniquely elicited a supramodal response starting at 100 ms after the oddball. Effective connectivity analyses identified shared interhemispheric interactions between inferior frontal gyri across modalities, as well as distinct modality-specific connectivity within early and associative sensory cortices. Our findings demonstrate that multi-feature prediction errors recruit both rapid supramodal integration mechanisms and hierarchically organized modality-specific pathways. These results advance our understanding of how the brain flexibly integrates multiple sensory expectation violations across different levels of cortical processing, providing new insights into the neural architecture supporting predictive perception.

**Author summary:** The brain constantly generates predictions about incoming sensory information. While many studies focus on simple violations of individual features, real-life events often involve simultaneous deviations across multiple sensory attributes. Our goal was to examine supramodal or modality-specific aspects of multi-feature prediction errors. Considering that the cortex needs to converge individual predictions from multiple pathways for multi-feature prediction, we hypothesised that multi-feature prediction errors rely on a mid-latency supramodal process in the inferior frontal cortex. In a high-density EEG study using a nested oddball paradigm, we examined neural responses to somatosensory and auditory stimulus deviations in one or two dimensions. Using temporal decoding, we revealed an early supramodal short-lived cortical process when double-deviants are detected. We also applied Parametric Empirical Bayes Modelling to show that multi-feature prediction errors not only rely on a common interhemispheric inhibition between inferior frontal gyri but also on various supramodal and modality-specific changes in effective connectivity across associative and modality-specific cortices.

## Introduction

According to the predictive coding theory of neural function, the brain continuously generates expectations about sensory input and signaling expectation violations as prediction errors (Clark, 2013; Egner & Summerfield, 2013; Friston, 2005). At each level of the cortical hierarchy, the brain compares top-down predictions with bottom-up sensory input to update its internal model of the environment. Prediction errors are propagated up the cortical hierarchy to iteratively minimize the discrepancy between expected and actual input (Friston, 2005; Mumford, 1992; Rao & Ballard, 1999). Prediction errors manifest as evoked neural responses to unexpected stimuli, and have been extensively studied using oddball paradigms in which single-deviant sequences of standard stimuli establish sensory expectations that are occasionally violated by deviant stimuli (Auksztulewicz et al., 2018; Garrido, Kilner, Kiebel, & Friston, 2009; R. Näätänen, Paavilainen, Rinne, & Alho, 2007; R. Näätänen, Simpson, & Loveless, 1982). Earlier studies found that neural responses encoding prediction errors can be elicited in a range of sensory modalities (Fardo et al., 2017; R. Näätänen et al., 2007; Ostwald et al., 2012; Pazo-Alvarez, Cadaveira, & Amenedo, 2003), and have modality-specific as well as supramodal components (Niedernhuber, Raimondo, Sitt, & Bekinschtein, 2022). Connectivity models of early mismatch responses in different sensory modalities consistently found a network spanning from bilateral primary sensory cortices to uni- or bilateral inferior frontal gyri via superior temporal gyri (Auksztulewicz & Friston, 2015; Chennu et al., 2016; Dietz, Friston, Mattingley, Roepstorff, & Garrido, 2014; Fardo et al., 2017; Garrido, Kilner, Stephan, & Friston, 2009; Ostwald et al., 2012; Holly N. Phillips et al., 2016; Timmermann et al., 2018). Most studies of predictive coding investigate neural responses to deviations of a single sensory feature (e.g., different auditory pitch (Garrido et al., 2008; R. Näätänen et al., 2007; H. N. Phillips, Blenkmann, Hughes, Bekinschtein, & Rowe, 2015), somatosensory two-point stimulation (Akatsuka, Wasaka, Nakata, Kida, & Kakigi, 2007)). Yet real-world events often involve simultaneous changes across multiple stimulus features, requiring the brain to process combined deviations within and across sensory modalities. One way to investigate this process is to study neural evoked responses to a double-deviant (e.g., pitch and location in audition, or intensity and site in touch) (Althen, Huotilainen, Grimm, & Escera, 2016; Gupta & Bhardwaj, 2022; Hansen, Højlund, Møller, Pearce, & Vuust, 2022; Ishida & Nittono, 2022; Levänen, Hari, McEvoy, & Sams, 1993; Paavilainen, Valppu, & Näätänen, 2001; Roach et al., 2020; Schroger, 1996; Schröger, 1995a; Takegata, Paavilainen, Näätänen, & Winkler, 1999; Takegata, Paavilainen, Näätänen, & Winkler, 2001; Wolff & Schröger, 2001). Previous work has shown that double-deviants elicit larger auditory local prediction errors than single-deviants, consistent with parallel predictive processing (Schröger, 1995b; Takegata et al., 1999). Due to its heightened sensitivity, the double-deviant prediction error has been shown to offer greater clinical utility in detecting cognitive impairments in neuropsychiatric populations than single-deviant responses Avissar et al. (2018). Although there is abundant evidence for supramodal aspects of prediction error encoding in the brain (Niedernhuber et al., 2022; Sabio-Albert, Fuentemilla, & Pérez-Bellido, 2025; Sanchez, Hartmann, Fuscà, Demarchi, & Weisz, 2020; Talsma, 2015), double-deviant prediction errors have mostly been studied within a sensory modality. As a result, it remains unclear whether prediction errors encoding more than one feature dimension (multi-feature prediction errors) rely on supramodal or modality-specific mechanisms.

Building on previous work in which we identified supramodal and modality-specific aspects of prediction error representations in the cortex using this dataset (Niedernhuber et al., 2022), we address whether there are supramodal or modality-specific representations which differentiate double-deviant from single-deviant prediction errors. Thirty participants performed somatosensory and auditory versions of a hierarchical oddball task while EEG was recorded (Bekinschtein et al., 2009). In this task, a local cortical prediction error is elicited when an unexpected stimulus is presented at the end of a group of spatiotemporally adjacent stimuli. When neural responses to deviant and standard stimuli are compared, deviant stimuli increase the amplitude of the Event-Related Potential (ERP) component overlapping with the N1–P2–N2 complex (R. Näätänen et al., 2007; Risto Näätänen, Tervaniemi, Sussman, Paavilainen, & Winkler, 2001). Although this response occurs in a similar time window as the classical auditory MMN, it does not necessarily exhibit the typical MMN morphology (i.e., a negative deflection) and often appears as a positive deflection. We therefore adopt the term ‘local effect’ in line with previous studies using this paradigm (Chennu et al., 2013, 2016). We introduced two types of prediction violations which differed from the standard stimulus in either one (single-deviant) or two (double-deviant) dimensions in each sensory modality (Bekinschtein et al., 2009; Niedernhuber et al., 2022). This task design choice enabled us to compare local neural responses elicited by double and single-deviants across modalities within the same paradigm. We reasoned that double-deviant violates predictions about the combination of features which requires the integration of multiple prediction channels that converge at higher-order association areas. Considering that the inferior frontal gyrus is consistently involved in local prediction error processing across modalities (Chennu et al., 2016; Fardo et al., 2017; Garrido, Kilner, Kiebel, et al., 2009; H. N. Phillips et al., 2015), we reasoned that double-deviant expectation violations might rely on a mid-latency supramodal process in the inferior frontal cortex, as well as activity in modality-specific pathways across the cortical hierarchy. Aligning with previous work (Schröger, 1995b; Takegata et al., 1999), we also hypothesised that double-deviant expectation violations will lead to a stronger cortical response than single-deviant ones. By combining Dynamic Causal Modelling (DCM) (David et al., 2006), Parametric Empirical Bayes (PEB) (Zeidman, Jafarian, Seghier, et al., 2019), and temporal decoding (King & Dehaene, 2014) at the same time, we were able to uncover shared and distinct cortical mechanisms and dynamics of multi-feature predictive coding between the somatosensory and auditory modality.

## Materials and methods

### Participants

30 individuals (15/15 female/male, aged 24.56 (+-4.52) mean+-STD) took part in the study. Only participants without hearing impairment and history of neurological or psychiatric disease were included in the study.

### Ethics

Participants freely consented to participate in the study in writing and were paid £10 per hour for a total duration of 3-3.5 h. The study was approved by the Cambridge Psychology Ethics Committee (CPREC 2014.25).

### Stimuli

The experimental design involved somatosensory and auditory stimuli (Figure 1). Somatosensory pressure stimuli applied with a custom-made device which applies mechanical pins to the fingertip with a Saia-Burgess 12.7mm stroke, 12v, 4W DC push-action solenoid with 0.3-0.6 N force, and no nominal delay after current onset) controlled by an Arduino Mega board. These stimuli were applied to the fingertip of either the left or the right hand. We generated auditory stimuli identical to (Bekinschtein et al., 2009; Chennu et al., 2013) by mixing three sinusoidal signals 500, 1000, and 2000 Hz for the single-deviant and standard stimulus or 350, 700, and 1400 Hz for the double-deviant stimulus in Matlab R2016. These were audibly presented to either the left or the right ear using EARTONE 3A insert earphones.

**Figure 1.**
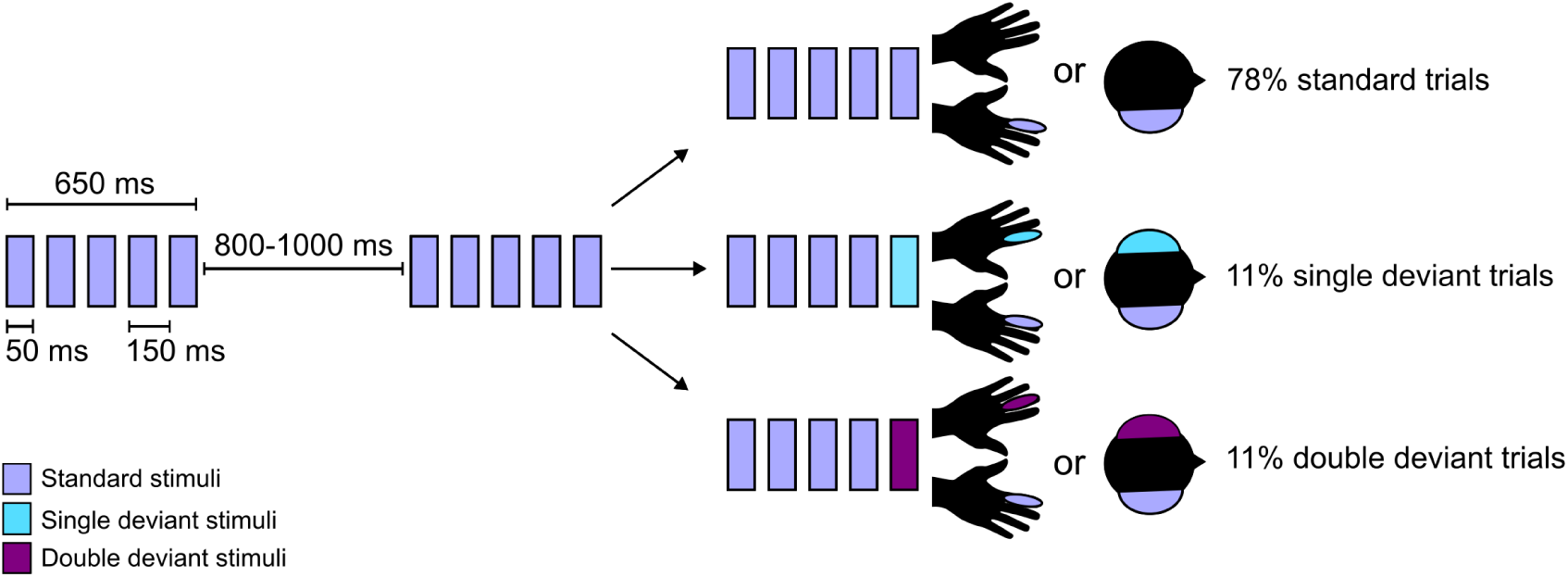
Experimental design. Stimulus sequences were presented in blocks of somatosensory or auditory stimulation. In a block, a stream of standard trials was occasionally interrupted by single-deviant or double-deviant trials. Each trial consisted of five stimuli with a duration of 50 ms and an interstimulus interval of 150 ms. Trials were separated by an intertrial interval of 800-1000 ms.

### Paradigm

This study uses a hierarchical oddball paradigm with nested local and global expectation violations in auditory and somatosensory blocks (Bekinschtein et al., 2009). For the present study, we focused exclusively on the local effects and analysed only standard and deviant trials matched for global predictability. Comparing neural responses to standard and deviant trials yields a short-lived prediction error response termed the local effect, which occurs within the same time window as other local prediction error responses such as the MMN or P3a (Bekinschtein et al., 2009).

Trials consisted of five stimuli presented with a stimulus onset interval of 150 ms and a duration of 50 ms. In deviant trials, four identical stimuli were followed by a deviant. We introduced two types of deviant trials to manipulate the strength of local violations: single-deviants differed from standards in one feature (laterality), while double-deviants differed in both laterality and stimulus type. Somatosensory standard trials consisted of five ipsilateral pressure stimuli delivered to the index finger of either hand. Single-deviant somatosensory trials involved stimulation of the index finger of the contralateral hand, and double-deviant trials involved stimulation of the middle finger of the contralateral hand.

Auditory standard trials consisted of five identical tones presented monaurally to one ear; single-deviant auditory trials involved a contralateral tone of the same pitch, while double-deviants involved a contralateral tone of a different pitch.

In both sensory modalities, the roles of standard and deviant stimuli were fixed throughout the experiment. Specifically, for auditory trials, one sound (e.g., 500 Hz) consistently served as the standard while another (e.g., 1000 Hz) served as the deviant; similarly, in the somatosensory condition, stimulation of one finger (e.g., left index) always functioned as the standard, and another (e.g., left middle) as the deviant. While this approach follows conventions in classical multi-feature oddball paradigms (Risto Näätänen, Pakarinen, Rinne, & Takegata, 2004), it does not implement full stimulus-role counterbalancing (e.g., AAAAB / BBBBA). As such, the resulting neural differences between standard and deviant conditions likely reflect both prediction-related responses and physical stimulus differences. Our current design therefore prioritizes ecological validity of sequential regularity but does not permit a dissociation of predictive and sensory-driven contributions to the observed effects.

For the present study, we focus on the local effect in a time window between 100-300 ms after onset of the last stimulus in a trial. In our dataset, streams of one trial type (globally standard trials) were interspersed with another (globally deviant trials). Single-deviants appeared either frequently or rarely within a block whereas double-deviants also deviated at a global level. Therefore, we included only deviant trials that were both locally and globally rare for both types of deviancy (and likewise for the standard trials). Each block contained 158–160 trials (∼4.5 min), with inter-trial intervals randomly sampled between 800–1000 ms in 50 ms steps. Blocks began with 15 globally standard trials to establish the frequent trial type, followed by 30–34 globally deviant trials interspersed within 112 globally standard trials. Globally deviant trials were equally likely to be preceded by 2–5 globally standard trials. To ensure that standards and deviants were matched for global predictability, only blocks where both trial types were globally consistent were included in the analysis (see (Niedernhuber et al., 2022) for analyses of other block types). This resulted in ∼60–68 deviant-standard trial pairs per condition.

The experiment always began with two somatosensory blocks to help participants fully grasp the task before proceeding. Piloting had shown that participants needed initial practice in the somatosensory condition to correctly identify and count deviant trials, and therefore starting with these blocks reduced confusion and experimental interruptions. White noise was played during somatosensory blocks to mask auditory cues from the tactile stimulator. Block order was pseudorandomized to avoid more than two consecutive blocks of the same modality. Two block types were presented twice for each modality and stimulus laterality (left/right).

Participants completed a handedness questionnaire (Oldfield, 1971), and tactile stimulation was calibrated to ensure identical perceived intensity across fingers before the experiment began. Auditory volume was individually adjusted for comfort. Participants were seated comfortably with visual fixation maintained. They were instructed to attend to the global structure of the stimulus sequences and to count rare patterns. After each block, participants reported the number of rare patterns detected. They were offered breaks between blocks.

### EEG preprocessing

We recorded 128-channel high density EEG data using a Net Amps 300 amplifier developed by Electrical Geodesics. Data were preprocessed in Matlab using the EEGLAB toolbox (Delorme & Makeig, 2004). Following the removal of channels on the neck, cheeks and forehead which record primarily artefacts resulting from head muscle and eye movements, we retained 92 channels for further analysis. We excluded the first 15 trials per block, which constitute the habituation phase in which a global prediction about incoming stimuli is generated, from further analysis. EEG data were filtered between 0.5 and 30 Hz and epoched in a time window between 200 ms before and 1300 ms after trial onset. Trial and channel artefacts were rejected by visual inspection of all channels. Noisy channels were excluded from further analysis and interpolated at a later stage. Artefacts originating from eye or muscle movements were visually identified and removed using independent component analysis (Delorme & Makeig, 2004). Following the interpolation of missing channels, data were re-referenced to the average and baseline-corrected relative to a 100 ms interval before the presentation of the last stimulus in the five-stimulus trial sequence.

### Cluster-based permutation of ERP voltage time series

We first demonstrated local effects in each sensory modality (auditory/somatosensory) and deviant type (single-/double-deviant) by comparing deviant-standard pairs. We also tested whether double-deviants elicit larger responses than single-deviants across modalities. To assess ERP differences, we used cluster-based permutation tests (Python v3.11.11, MNE v1.9.0; (Gramfort et al., 2013; Maris & Oostenveld, 2007)), a non-parametric approach that controls for multiple comparisons in spatiotemporal EEG data. Trial numbers were equalized by subsampling. Analyses were restricted to the 100–300 ms window typical for MMN responses (R. Näätänen, Tervaniemi, Sussman, Paavilainen, & Winkler, 2001). Clusters were formed from spatiotemporally adjacent t-values and only retained if they contained at least five electrodes and p < .05. Cluster-level significance was determined using a two-tailed *Monte Carlo* procedure with 2000 permutations (*α* = 0.05).

### Temporal decoding

We employed temporal decoding (also known as the temporal generalisation method) to assess commonalities and differences in the cortical representations of double-deviants and single-deviants within and between sensory modalities ((King & Dehaene, 2014)). Temporal decoding is a machine learning approach used to characterise how cognitive operations unfold in time (King & Dehaene, 2014). Classifiers are trained on data from a specific time point and tested across all time points. This generates a temporal generalisation matrix which reveals how neural information is maintained in the cortex (King, Gramfort, Schurger, Naccache, & Dehaene, 2014). Diagonal elements reflect classification performance when training and testing occur at the same time point, while off-diagonal elements reflect the extent to which neural representations generalize across time. Although this method provides insight into the temporal dynamics of cortical processes using EEG sensor data, it does not allow inferences about spatial sources.

Classification was implemented in Python (v3.11.11) using MNE (v1.9.0) (Gramfort et al., 2013). To prevent class imbalance, trial numbers were equalised across conditions by randomly deselecting trials from conditions with more trials. Logistic regression models were trained on normalised data. For within-condition decoding, model training and evaluation used stratified 5-fold cross-validation. For comparisons between conditions, classifiers were trained on one dataset and tested on independent datasets from other conditions. When comparing single- and double-deviants directly, standard trials were partitioned into disjoint subsets to ensure independence between training and testing sets. Classification was performed within a 600 ms window following the onset of the last stimulus in a trial, and decoding performance was evaluated using the Area under the Curve-Receiver Operating Characteristic (AUC-ROC). Statistical significance was assessed via Monte Carlo cluster-based permutation tests with 4096 random partitions and two-tailed paired t-tests (p < .05) (Maris & Oostenveld, 2007).

This pipeline was applied to: (1) auditory single-deviant vs. standard, (2) auditory double-deviant vs. standard, (3) somatosensory single-deviant vs. standard, and (4) somatosensory double-deviant vs. standard trials. Additionally, classifiers trained on one condition (e.g., auditory single-deviant vs. standard) were tested across other conditions to assess generalisation across deviance levels and modalities. Classifiers were also trained and tested to directly discriminate single-deviant vs. double-deviant trials within and across modalities.

### Dynamic Causal Modelling and Parametric Empirical Bayes

Based on previous work (Zeidman, Jafarian, Seghier, et al., 2019), we used DCM and PEB to examine commonalities and differences in effective connectivity changes due to double-deviants between the somatosensory and auditory modality. DCM for Event-Related Potentials (ERPs) is a validated, neurophysiologically grounded Bayesian source reconstruction method used to describe causal interactions between cortical sources elicited by ERPs (David et al., 2006). A set of differential equations describe effective connectivity, which is the rate of change at which a neuronal population in a cortical source exerts influence over another, and the modulation of effective connectivity elicited by an experimental intervention (Friston & Kiebel, 2009; Stephan et al., 2010). Based on a framework for neural mass models developed by Jansen and Rit (Jansen & Rit, 1995), DCM for ERPs relies on a neural mass model which describes neuronal activity in three cortical layers: an excitatory pyramidal cell population in the granular layer, an inhibitory interneuron population in the supra-granular layer and a subpopulation of deep pyramidal cells in the infra-granular layer. Interactions between these neuronal subpopulations are described via extrinsic connections which describe information flow between sources traversing white matter and intrinsic connections which represent coupling within a cortical source. The theoretical framework underpinning DCM assumes a hierarchical organisation of cortical areas. Based on a proposal by Felleman and Van Essen (Felleman & Van Essen, 1991), connections between sources are directional and can either be forward, backward or lateral. Earlier work characterising connections in the visual cortical hierarchy (Van Essen & Maunsell, 1983) frame pathways in the cortex as either ascending or forward (originating in a lower-ranked cortical area and terminating in a higher-order cortical area), descending or backward (originating in a higher-order cortical area and terminating in a lower-order area), or lateral connections between two cortical areas at the same hierarchical level. DCM is extended by a spatial forward model which translates cortical activity arising from depolarised pyramidal cells to responses from EEG sensors described as an equivalent current dipole (ECD) model. Taken together, this renders DCM a spatiotemporal generative model of context-related changes in effective connectivity between cortical sources. To perform group-level analyses, we can construct a PEB model over the resulting DCM parameters (Zeidman, Jafarian, Seghier, et al., 2019). (The Bayesian Model Average of) PEBs can be described as a General Linear Model:

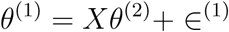

*θ*^(1)^ refers to the connection weights for each participant. *Xθ*^(2)^ represent the estimated influence of each covariate on each connection entered into the model. A design matrix *X* describes the regressors which model the impact of each covariate on each connection. ∈^(1)^ adds an error term which models random effects variability between participants. PEB models can be used to describe commonalities and differences in DCM parameters between groups (Zeidman, Jafarian, Corbin, et al., 2019; Zeidman, Jafarian, Seghier, et al., 2019).

For each sensory modality (auditory/somatosensory) and deviance level (single-deviant/double-deviant), we used DCM for ERPs to estimate effective connections elicited in a network recruiting primary and secondary somatosensory cortices (S1, S2), primary auditory cortices (A1), both superior temporal gyri (STG) and both inferior frontal gyri (IFG). We selected bilateral A1, STG, IFG, S1, and S2 based on previous dynamic causal modeling studies of auditory and somatosensory mismatch responses (Chennu et al., 2016; Fardo et al., 2017; Garrido, Kilner, Kiebel, et al., 2009; H. N. Phillips et al., 2015). To investigate how both contralateral deviant and ipsilateral standard inputs jointly shape prediction error signals, we modeled driving inputs bilaterally to the corresponding primary sensory cortices (A1 for auditory trials, S1 for somatosensory trials). For somatosensory trials, we modeled driving inputs to bilateral S1 to accommodate potential early bilateral activation and interhemispheric interactions (e.g., (Tamè, Braun, Holmes, Farnè, & Pavani, 2016)), and to maintain symmetry with the auditory models. While somatosensory afferents are primarily contralateral, this modeling choice avoids overly constraining the input structure and allows the group-level analysis to prune unsupported parameters via empirical Bayes optimization (Friston et al., 2016).

A1 corresponds to the primary auditory cortex (posteromedial Heschl’s gyrus), while STG encompasses higher-order auditory areas, including regions corresponding to A2. Similarly, S1 and S2 represent primary and secondary somatosensory cortices respectively, commonly involved in tactile deviance detection (Chennu et al., 2016; Fardo et al., 2017; Garrido, Kilner, Kiebel, et al., 2009; H. N. Phillips et al., 2015).

We used MNI coordinates provided in (Garrido et al., 2008) as dipole locations for the primary auditory cortex, superior temporal gyrus and inferior frontal gyrus for the forward projection from source to sensor space. To identify sources for the primary and secondary somatosensory cortex, we performed an auxiliary source reconstruction on somatosensory deviant trials using Brainstorm (Tadel, Baillet, Mosher, Pantazis, & Leahy, 2011). We applied the symmetric Boundary Element Method available from the open-source software OpenMEEG to estimate a forward model using the cortex surface as a source space. Since individual MRIs were unavailable, we used the default anatomical template MRI ICBM152_2019 in Brainstorm. We removed the DC offset and estimated the noise covariance using minimum norm estimators using data from baseline activity in a 100 ms time window before the onset of the first stimulus in a trial. In a further step, we used minimum norm estimation which constrains sources to be perpendicular to the cortex surface with a current density map to obtain an inverse solution. MNI coordinates for sources were obtained using visual inspection (Supplementary Figure 1).

A model was fitted to each deviant-standard contrast within 0-250 ms after onset of the last stimulus in a trial per participant. For each sensory modality and deviance level, we modelled changes in effective connectivity due to a deviant stimulus in a network with auditory, somatosensory and frontal sources. This resulted in 30 DCMs for each of the four conditions in our 2×2 design (deviance: single/double, sensory modality: somatosensory/auditory). Trial numbers in each condition pair were equalised and no Hanning window was used. Having been furnished with a forward model, the model was inverted for each participant. Model inversion approximates the posterior probability of connectivity parameters by minimising a free-energy bound on the log-evidence (Friston, Mattout, Trujillo-Barreto, Ashburner, & Penny, 2007) and serves to constrain interactions between nodes to cortical sources (David et al., 2006).

We assessed commonalities and differences in effective connectivity elicited by multi-feature prediction errors in the somatosensory and auditory modality using a PEB-of-PEBs approach. We identified the most likely modulation of connections by inverting a DCM including all plausible connections between auditory, somatosensory and frontal sources for each participant and for each contrast type (using spm_dcm_erp.m)(Zeidman, Jafarian, Corbin, et al., 2019; Zeidman, Jafarian, Seghier, et al., 2019). We proceeded to construct a PEB over DCM parameters modelling commonalities and differences in effective connections between double-deviant and single-deviant stimuli (B-Matrix) for each sensory modality (using spm_dcm_peb.m). Each PEB included the group mean of each connectivity parameter and the effect of deviance on the group mean as regressors. This procedure outputs a somatosensory and an auditory PEB encoding the group mean and the effect of deviance for each connection. Finally, we examined commonalities and differences in effective connectivity between sensory modalities. For that, we constructed a PEB over somatosensory and auditory PEB parameters using the mean of all participants and the difference between groups as regressors. We performed Bayesian Model Reduction of posterior densities over PEB parameters to remove connections which do not contribute to the model evidence using a greedy search algorithm (using spm_dcm_peb_bmc.m). Having pruned redundant connections, we carried out Bayesian Model Averaging across the 256 models with the best fit identified by the search. The resulting PEB encodes parameters for the mean connectivity over participants, the main effect of sensory modality and deviance as well as their interaction for each connection. We retained connections with a posterior probability of at least 95% which can be interpreted as strong evidence.

## Results

### Double-deviants produce stronger cortical responses regardless of sensory modality

In the first step shown in Figure 2, we identified cortical responses to double-deviant and single-deviant stimuli using cluster-based permutation tests of ERP voltage time courses in a time window between 100-200 ms. We established differences between deviant and standard trials for auditory single-deviants (cluster t = −0.135 ± 2.260, p = 0.001), double-deviants (cluster t = 0.092 ± 4.594, p < 0.001), as well as somatosensory single-deviants (cluster t = −0.033 ± 1.550, p = 0.009) and double-deviants (cluster t = 0.008 ± 3.166, p < 0.001). In line with previous work (Auksztulewicz & Friston, 2015), we observed that double-deviants produce a stronger ERP response amplitude than single-deviants in the auditory (cluster t = 0.122 ± 4.688, p < 0.001), and somatosensory modality (cluster t = 0.056 ± 2.948, p < 0.001)

**Figure 2.**
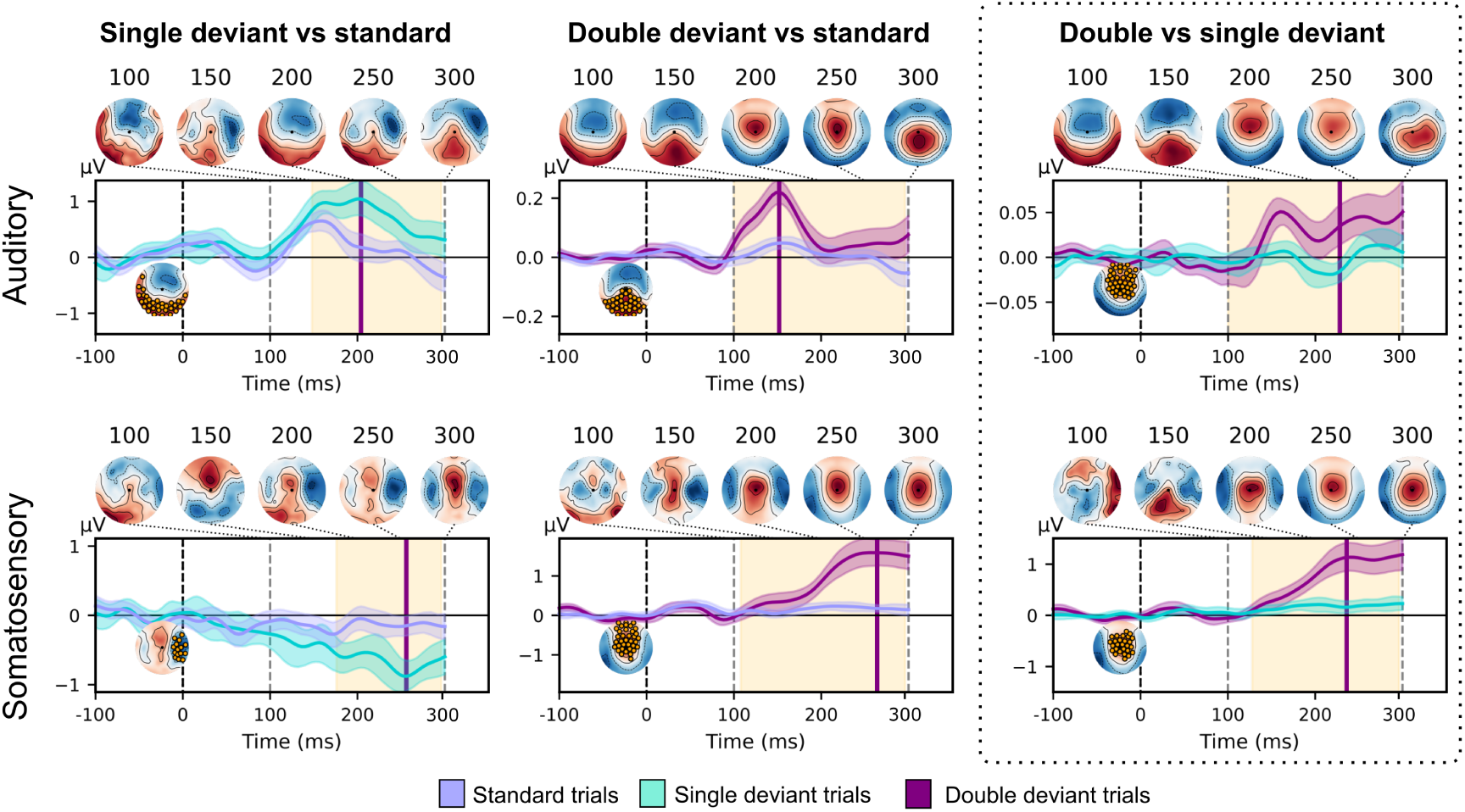
ERP responses to single and double deviants across sensory modalities. Results from a cluster-based permutation test on ERP time courses reveal a difference between deviant and standard trials for somatosensory and auditory single-deviants and double-deviants. A comparison of single-deviants and double-deviants showed a stronger ERP response to double-deviants in both sensory modalities. For each comparison, mean ERP voltage amplitude time courses are shown (with the 95% confidence band shaded). 0 denotes the onset of the fifth stimulus in a trial. Time segments in which ERP time courses differ are highlighted in orange. In the ERP voltage time course plot, the time point at which the difference between both conditions is maximal is marked with a purple line. In the time course plot, we show the topography corresponding to the maximal difference between conditions. The cluster with the highest summarised t-value and *p* < .05 is shown in orange. Topographies of the difference between conditions are shown at 100, 200 and 300 ms

### Transient supramodal activity encodes multi-feature prediction errors

Our goal was to assess whether cortical networks converging prediction errors from different pathways are supramodal or modality-specific. Based on the theory that frontoparietal activity is crucial to encode this difference across sensory modalities, we hypothesised that commonalities and differences between single-deviants and double-deviants might emerge in a mid-latency time-window. Building on previous studies showing that sensory deviants elicit activity in supramodal brain areas (Covington & Polich, 1996; Niedernhuber et al., 2022; Walz et al., 2013), we uncovered supramodal and modality-specific cortical dynamics specific to double-deviants using temporal decoding (Figures 3 and 4).

**Figure 3.**
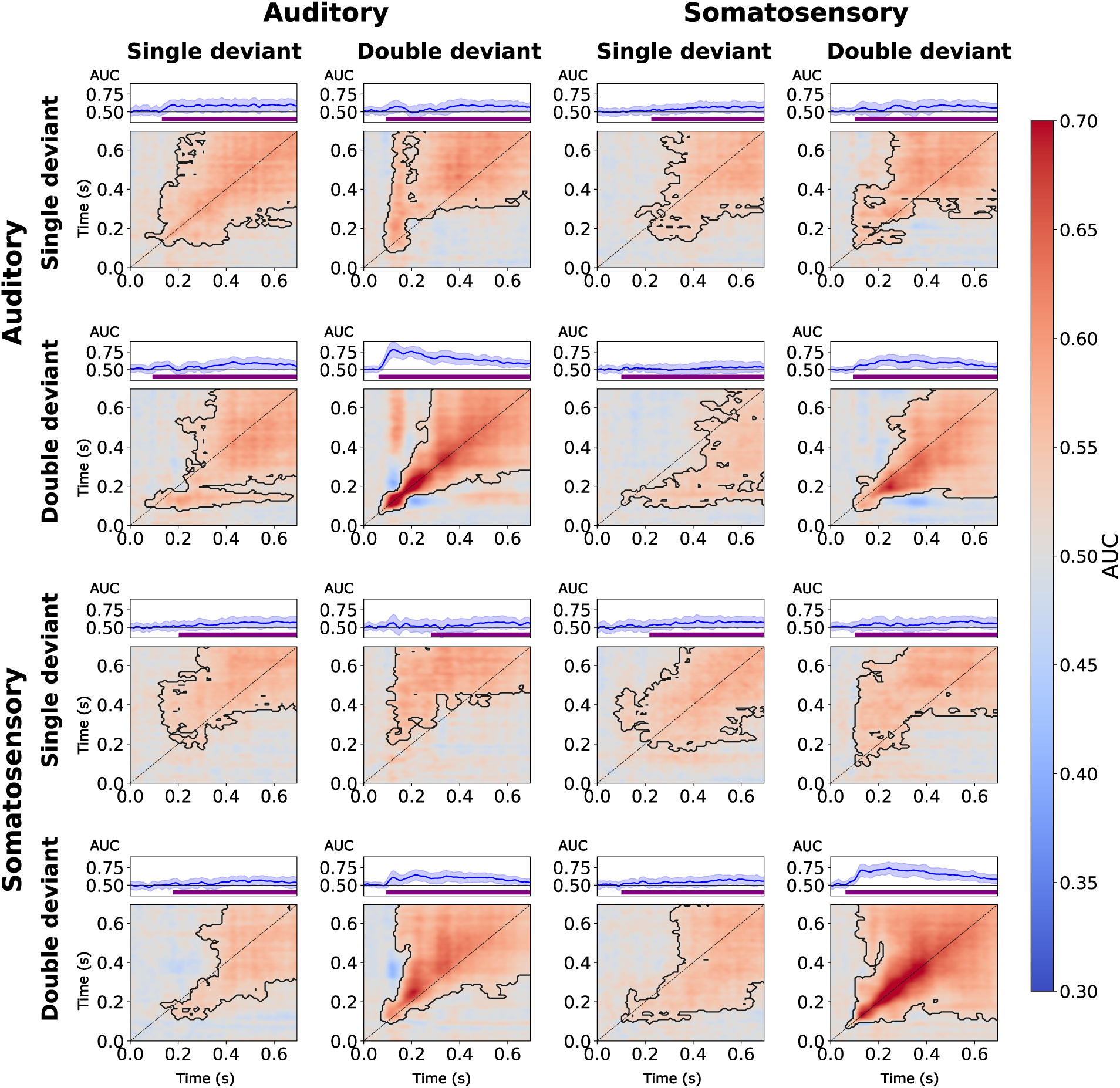
Temporal decoding analysis contrasting double-deviants and single-deviants within and across sensory modalities. AUC-ROC classification score matrices are plotted on a red-to-blue gradient. Adjacent classification score clusters which differ from chance were identified in a cluster-based permutation test and delineated using a dotted line. The left top panel shows the average diagonal classification performance. Each adjacent time course represents mean scores (and the standard deviation shaded) obtained from a classifier tested at a target time point (100, 200, 300 ms) and trained on all remaining time points. Purple bars highlight significant differences between single-deviant and double-deviant classification score series with *α* = 0.05 (B).

**Figure 4.**
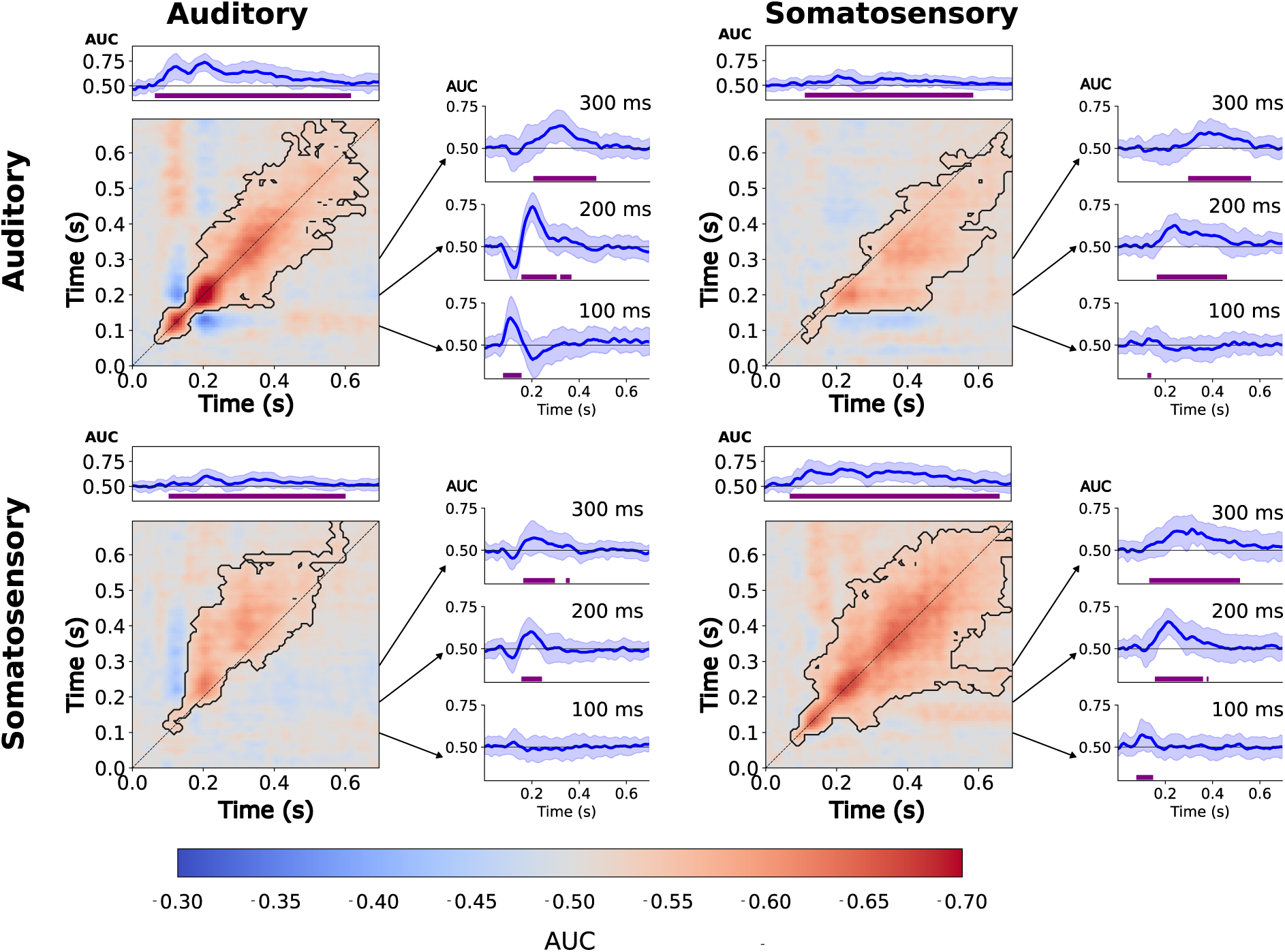
Temporal decoding analysis comparing deviant-standard pairs within and across sensory modalities. Classification performance is shown as AUC-ROC score matrices, visualized using a red-to-blue gradient. Significant classification clusters (p < 0.05) were identified using cluster-based permutation tests and are marked with black outlines. In each matrix, the top panel displays mean diagonal classification accuracy (and the standard deviation shaded). Purple bars indicate time intervals with significant differences between single-deviant and double-deviant classification series (*α* = 0.05, B).

#### Single-deviants

In line with previous work (King et al., 2014; Niedernhuber et al., 2022), classifiers trained to distinguish single-deviant and standard trials generalised from ∼200 ms after oddball onset until the end of the trial regardless of contrast. Sustained decoding along both diagonal and off-diagonal elements suggests that the neural representations supporting deviance detection remain active or are re-engaged over several hundred milliseconds. We observed this temporal generalisation pattern for both the auditory (cluster t = 2.899 ± 2.214, p < 0.001) and somatosensory single-deviant (cluster t = 2.276 ± 2.077, p < 0.001). Corroborating past work (Niedernhuber et al., 2022), cross-decoding single-deviants between sensory modalities revealed a P3b-like supramodal activity pattern with some temporal shifts. For single-deviants, we replicated this finding for somatosensory-to-auditory cross-decoding (cluster t = 1.905 ± 1.904, p < 0.001) and vice versa (cluster t = 1.758 ± 1.995, p < 0.001).

#### Double-deviants

Auditory double-deviants could be decoded along the diagonal between ∼100–200 ms and only generalised across the remaining time window from ∼200 ms (cluster t = 4.180 ± 4.446, p < 0.001). Based on (King et al., 2014), we speculate that this pattern might reflect a serial cortical process in a mid-latency time window that transitions into a more sustained activation profile (King et al., 2014). For its somatosensory counterpart, we observed temporal generalisation from ∼100 ms until trial end (cluster t = 5.198 ± 3.927, p < 0.001), suggesting that cortical activity is maintained in a single cortical network. When classifiers were trained and tested to discriminate double-deviant and standard trials across sensory modalities, we uncovered extensive temporal generalisation pattern from ∼200 ms with some temporal shifts. This was observed when crossdecoding from the auditory to the somatosensory modality (cluster t = 2.695 ± 3.566, p < 0.001) and vice versa (cluster t = 3.346 ± 3.615, p < 0.001).

#### Supramodal activation patterns support single-deviants and double-deviants

Single-deviants and double-deviants were found to share a common late activity pattern between sensory modalities. This pattern was found when classifiers were trained using the auditory single-deviant contrast and tested on the somatosensory double-deviant contrasts (cluster t = 2.196 ± 2.382, p < 0.001) and vice versa (cluster t = 1.063 ± 1.670, p < 0.001). We found a similar pattern when crossdecoding between the somatosensory single-deviant and auditory double-deviant, (cluster t = 1.883 ± 1.819, p < 0.001) and the somatosensory double-deviant and single-deviant (cluster t = 1.357 ± 2.202, p < 0.001). In sum, our findings support the notion that there is a higher-order supramodal network supporting multi-feature and single-feature predictive coding (Corbetta & Shulman, 2002; Macaluso, Frith, & Driver, 2002; Niedernhuber et al., 2022).

#### Modality-specific activity differentiates single-deviants and double-deviants

Finally, we tested classifiers to distinguish single-deviants and double-deviants (Figure 4). Our results show that there is modality-specific activity over several hundred millisecond, but cortical activation was only briefly maintained. This was found for the somatosensory (cluster t = 2.382 ± 2.451, p < 0.001) and auditory contrast (cluster t = 1.433 ± 2.474, p < 0.001). Interestingly, cross-decoding from the auditory to the somatosensory modality revealed a similar activation pattern (cluster t = 1.083 ± 2.127, p < 0.001), and vice versa (cluster t = 0.842 ± 1.979, p < 0.001).

In sum, both single-deviants and double-deviants lead to a late temporal generalisation pattern, but only double-deviants were found to elicit a supramodal activity pattern briefly maintained along the diagonal. Since we only focus on trials in which local deviants also deviate in a block, and the decodability of the P3b-like global response is very strong relative to the MMN-like local response, the shape of the local response differs from previous work (King et al., 2014; Niedernhuber et al., 2022). Importantly, our findings suggests that these shared short-lived activation patterns are specific to the double-deviant. Whereas both single-deviants and double-deviants elicit a late supramodal activation pattern, only responses to double-deviants were supported by a supramodal transient process (Figure 4). With these results, we replicated and extended earlier findings that deviants of sensory stimulus groups elicit a P3b-like difference wave from ∼200 ms until trial end in either sensory modality (King et al., 2014; Niedernhuber et al., 2022). We also provide corroborating evidence for our own finding that shared representations between the auditory and the somatosensory modality emerge from ∼200 ms for both the double-deviant and single-deviant (Niedernhuber et al., 2022).

### Supramodal and modality-specific effective connectivity underpins multi-feature prediction errors

Having identified a transient supramodal process for multi-feature prediction errors, we expected that multi-feature prediction errors might be encoded in supramodal and modality-specific connectivity changes in the frontoparietal cortex. Subsequently, we describe results from a DCM-PEB analysis used to assess commonalities and differences in connectivity across the somatosensory and auditory cortical hierarchy. We designed a DCM with all plausible bidirectional connections between primary and secondary sensory as well as frontal cortices in the somatosensory and auditory hierarchy (Figure 5).

**Figure 5.**
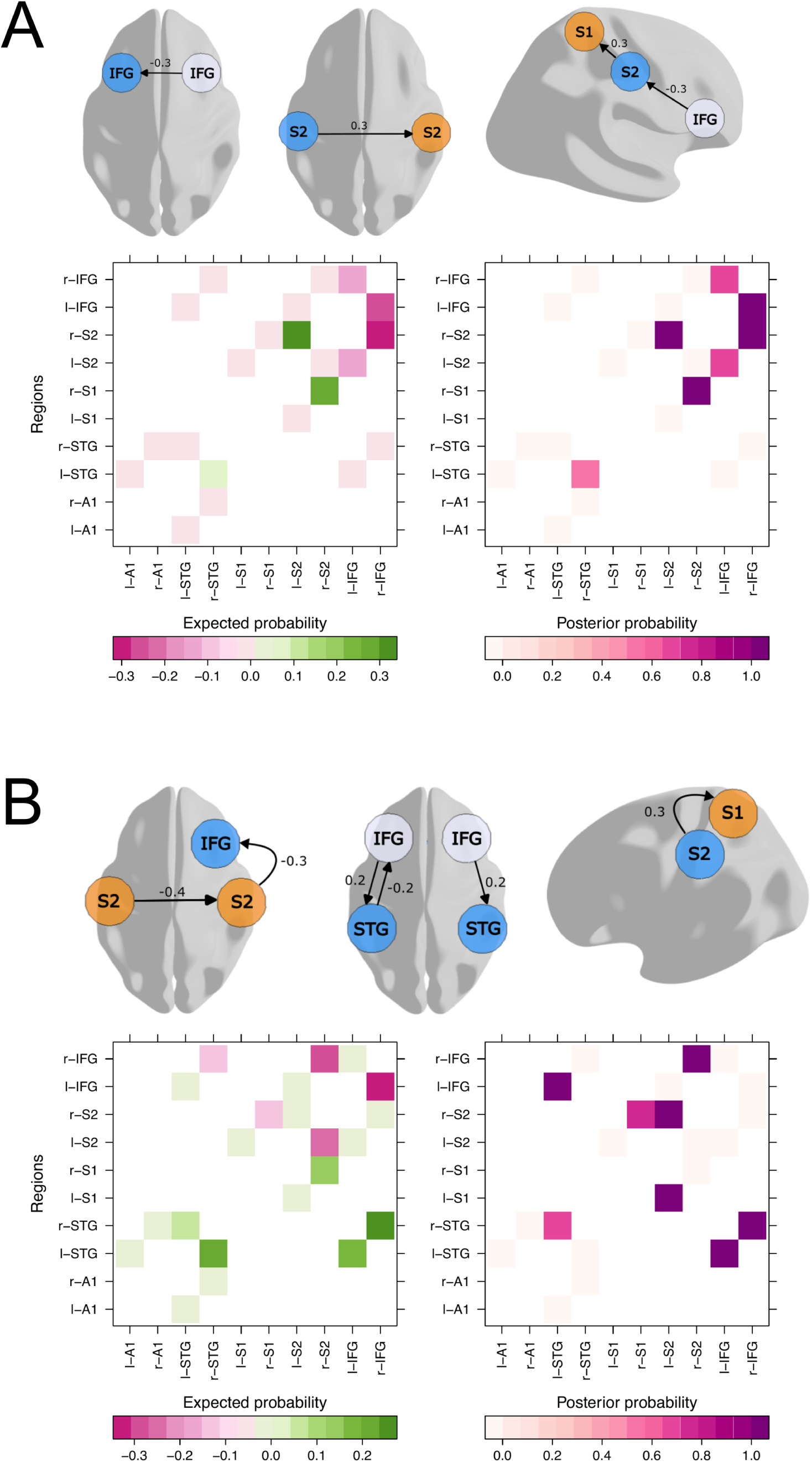
Effective connectivity modelling. The top panel shows commonalities (A) and differences (B) in effective connectivity between single and double-deviants. Effective

Our analysis revealed commonalities and differences in multi-feature predictive coding at early and late stages of the somatosensory and auditory hierarchy. Differences between sensory modalities became evident across associative somatosensory and frontal cortices. As shown in Figure 5, B), multi-feature prediction errors manifested in decreased connectivity between the right secondary somatosensory cortex and inferior frontal gyrus as well as interhemispheric connectivity between secondary somatosensory cortices in the somatosensory relative to the auditory condition. Results from a comparison of single-deviants and double-deviants for each sensory modality are shown in Supplementary Figure 2. Double-deviants modulated connectivity from the left secondary to primary somatosensory cortex more strongly in the somatosensory modality. Conversely, connectivity changes from the inferior frontal gyrus to the superior temporal cortex elicited by double-deviants were reduced in the auditory modality in each hemisphere. We also discovered commonalities in the processing of multi-feature prediction errors between sensory modalities (Figure 5, A). We identified a common inhibition in an interhemispheric connection between inferior frontal gyri. Interestingly, we also found commonalities in information flow between frontal, associative and early sensory cortices. Auditory and somatosensory networks processing multi-feature prediction errors share an excitatory connection descending from the right inferior frontal gyrus to the ipsilateral secondary somatosensory cortex and between secondary somatosensory cortices. We also identified a common excitatory input from the right secondary to the right primary somatosensory cortex. In sum, supramodal connectivity encoding multi-feature prediction errors can be identified between lower, intermediate and higher levels of the cortical hierarchy. Our results complement earlier findings demonstrating direct interactions between early and associative auditory and somatosensory cortices (Pérez-Bellido, Anne Barnes, Crommett, & Yau, 2018; Ro, Ellmore, & Beauchamp, 2013). Our results support the view that information processing in the cortex is multisensory in early and late stages of sensory processing (Ghazanfar & Schroeder, 2006).

## Discussion

Our study examined to what extent multi-feature prediction errors are supported by supramodal or modality-specific networks. We made three key observations: First, we identified a difference in voltage amplitude time courses between single-deviants and double-deviants in each sensory modality. Second, we showed that both double-deviants and single-deviants elicit supramodal activity over an extended late time window using temporal decoding. However, double-deviants were found to rely on a specific mid-latency supramodal process. Lastly, we performed a follow-up comparison of connectivity changes linked to multi-feature prediction errors between both sensory modalities. Using DCM-PEB, we identified common connections supporting multi-feature prediction errors between sensory modalities not only between bilateral inferior frontal cortices but also in lower-order sensory cortices. We also identified differences in connectivity between single-deviants and double-deviants common between sensory modalities. Overall, we identified a supramodal process for the detection of double-deviants which manifests in shared cortical dynamics and frontal and modality-specific information flow. Beyond that, we demonstrate that supramodal and modality-specific mechanisms supporting multi-feature prediction errors are not limited to the inferior frontal cortex but ubiquitous across the cortical hierarchy (Ghazanfar & Schroeder, 2006; Schroeder & Foxe, 2005).

In agreement with our results, the inferior frontal gyrus has been found to selectively respond to rare deviant stimuli in the visual (Hampshire, Thompson, Duncan, & Owen, 2009; Kirino, Belger, Goldman-Rakic, & McCarthy, 2000), auditory (Auksztulewicz & Friston, 2015; Chennu et al., 2016; Garrido et al., 2008; H. N. Phillips et al., 2015; Holly N. Phillips et al., 2016), and somatosensory modality (Allen et al., 2016; Fardo et al., 2017; Ostwald et al., 2012). Involvement of the inferior frontal cortex was consistently found in various sensory oddball paradigms, positioned on top of a hierarchy of modality-specific cortices (Chennu et al., 2016; Corbetta, Patel, & Shulman, 2008; Downar, Crawley, Mikulis, & Davis, 2000; Fardo et al., 2017; Garrido, Kilner, Stephan, et al., 2009; Kirino et al., 2000; Ostwald et al., 2012). Beyond past studies, our finding provides direct evidence that multi-feature predictive coding relies on a supramodal inferior frontal inhibition between hemispheres. Unexpectedly, we identified supramodal connections between secondary somatosensory cortices and even between the primary and secondary somatosensory cortex. Although not included in our prediction, supramodal aspects of multi-feature predictive coding in early processing stages are plausible. There is rich evidence that multisensory information is processed across the cortex (Ghazanfar & Schroeder, 2006). Indeed, the secondary somatosensory cortex borders the temporoparietal junction and the auditory cortex where patches of somatosensory but also multisensory neurons have been found (Brett-Green, Fifková, Larue, Winer, & Barth, 2003; Menzel & Barth, 2005). White-matter projections between the secondary somatosensory cortex and auditory cortex have been documented in the literature (Ro et al., 2013). In addition to audiotactile integration, the somatosensory cortex has been found to respond to purely auditory inputs (Pérez-Bellido et al., 2018). In sum, our findings support a role for the inferior frontal cortex as a key region for multi-feature predictive coding alongside supramodal and modality-specific activity in earlier cortical regions.

A possible explanation for our findings is that double-deviants are more salient and attract more stimulus-driven attention than single-deviants via a bottom-up mechanism (Parr & Friston, 2019). Canonically, a two-stream system is thought to shift attention to novel stimuli. A ventral frontoparietal attention network deploys exogenous bottom-up salience to double-deviant sensory events from the bottom up (Hampshire, Chamberlain, Monti, Duncan, & Owen, 2010). Conversely, a dorsal attention network binds endogenous goal-driven attention to sensory events based on top-down expectations (Corbetta et al., 2008; Corbetta & Shulman, 2002). Both systems are co-activated when relevant sensory deviant stimuli are detected, regardless of which sensory modality is stimulated (Corbetta et al., 2008; Downar et al., 2000). There is evidence for supramodal (Banerjee, Snyder, Molholm, & Foxe, 2011; Downar et al., 2000; Farah, Wong, Monheit, & Morrow, 1989; Golay, Hauert, Greber, Schnider, & Ptak, 2005; Green, Doesburg, Ward, & McDonald, 2011; Macaluso, Eimer, Frith, & Driver, 2003; Macaluso et al., 2002; Spence & Driver, 1997) and modality-specific (Chambers, Stokes, & Mattingley, 2004; Fleming, Njoroge, Noyce, Perrachione, & Shinn-Cunningham, 2024; Michalka, Kong, Rosen, Shinn-Cunningham, & Somers, 2015; Abigail L. Noyce, Cestero, Michalka, Shinn-Cunningham, & Somers, 2017; Abigail L. Noyce et al., 2022; Santangelo, Fagioli, & Macaluso, 2010) aspects of orienting attention toward a target. These studies have shown that interdigitated regions in lateral frontal cortex preferentially support visual or auditory attention, yet also exhibit flexible recruitment depending on task demands (such as the spatial or temporal nature of the information) which reflects a mixture of modality-specific and supramodal characteristics. As a part of the ventral stream, the inferior frontal gyrus has been thought to support attention reorientation to relevant stimuli (Corbetta & Shulman, 2002). It is therefore possible that signatures of multi-feature prediction errors might share supramodal information flow between inferior frontal gyri due to salience-induced attentional shifts. Supporting the notion that attention biases activity at early and late stages of the cortical hierarchy (Dugué, Merriam, Heeger, & Carrasco, 2020; Hopfinger & West, 2006; Saenz, Buracas, & Boynton, 2002), attention might also explain supramodal and modality-specific information flow in primary and secondary cortices.

Although our design did not explicitly manipulate attention, we acknowledge that both stimulus-driven and top-down attentional processes might contribute to the observed neural dynamics. Participants attended to all deviant stimuli, regardless of deviancy or modality. While top-down task demands might have contributed to supramodal decoding performance, these demands were equivalent across conditions and therefore cannot account for differences between deviancy levels or sensory modalities. Working memory demands might also have contributed to the late supramodal activation pattern observed in the temporal decoding results, which resembles a P3b-like response—an ERP component typically linked to attention and memory processes (King et al., 2014). Previous research, including our own work (Niedernhuber et al., 2022; Walz et al., 2013), has shown that the P3b recruits a supramodal network. Interestingly, while both single-deviant and double-deviants evoked this extended activation, only double-deviants activated a supramodal transient response. This suggests that early stages of sensory prediction might involve a supramodal mechanism for multi-feature predictive coding, whereas later responses might reflect more general task-related cognitive processes, such as attention and memory load.

### Limitations

While our findings provide new insights into the neural encoding of multi-feature prediction errors, several limitations should be noted. First, our study did not include behavioral measures directly tied to prediction error strength or perceptual salience, limiting our ability to link neural responses to subjective experience or task performance. Second, although we employed temporal decoding and dynamic causal modeling to characterize neural dynamics and connectivity, EEG inherently offers limited spatial resolution, and precise source localization, particularly in deep or overlapping cortical regions, remains approximate and tied to the parameters and decisions of the inverse solution of the DCM. Individual MRI scans were unavailable, which reduces the spatial accuracy of DCM-PEB. Third, our paradigm was not designed to dissociate prediction error from its potential downstream effects, such as attentional shifts or memory encoding. As a result, interpretations involving higher-order cognitive processes should remain cautious and speculative. Future studies combining neuroimaging modalities or incorporating behavioral readouts could help address these open questions. Another concern is that our current work might not constitute a substantial advance relative to another paper published by our group (Niedernhuber et al., 2022). In that publication (Niedernhuber et al., 2022), we focused on identifying supramodal neural correlates of local and global prediction errors using temporal decoding methods. The current study re-analyses this dataset with a focus on the distinction between single-deviant and double-deviants. This contrast was not addressed in the previous paper. In addition, we include new analyses of effective connectivity using DCM-PEB, which were not part of the earlier work.

While our paradigm was designed to elicit local prediction errors through violations of simple sensory regularities, we acknowledge that the physical features of deviant stimuli, particularly in the double-deviant condition, differ from those of standard stimuli in both identity and laterality. As a result, the observed neural responses likely reflect a combination of expectation-based and sensory-driven processes. Crucially, this is consistent with predictive coding models, which posit that prediction errors emerge when sensory input diverges from an internal model, irrespective of whether the deviation involves identical or physically distinct stimuli (Friston, 2005; Garrido, Kilner, Stephan, et al., 2009). In this view, the key determinant for predictive coding is not the physical sameness of the stimulus, but the violation of predicted sensory regularities. This interpretation aligns with prior work showing that MMN and related responses can reflect hierarchical prediction errors even when elicited by physically distinct inputs (Chennu et al., 2016; Wacongne et al., 2011). Nonetheless, because stimulus roles were not counterbalanced in our design, we cannot isolate the relative contributions of sensory-specific features and “pure” or cognitive predictive processes. Future studies could improve interpretability by systematically counterbalancing stimulus identities across standard and deviant roles to more precisely dissociate these components.

## Data and code availability

Data can be found here: https://zenodo.org/records/15330669. Code for the study can be found here: https://osf.io/e8hps/.

## Author contributions

Maria Niedernhuber: conceived and designed the study; collected the data; analysed the data; interpreted the results; wrote the manuscript. Micah Allen: conceived and designed the study (conceptual framework, analysis); interpreted the results; edited the manuscript. Francesca Fardo conceived and designed the study (conceptual framework, analysis); interpreted the results; edited the manuscript. Tristan Bekinschtein conceived and designed the study (conceptual framework, analysis); interpreted the results; edited the manuscript; acquired funding for the study.

