## Supplementary for "Supramodal and modality-specific neural information supports multi-feature prediction errors across cortical levels"

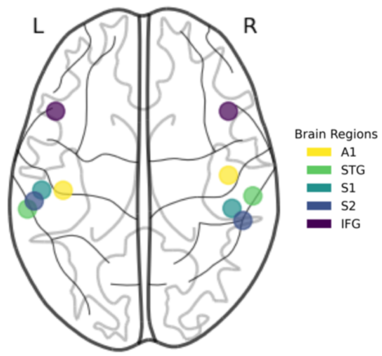


| Source | Left | Right |
|---|---|---|
| Primary auditory cortex (A1) | -42, -22, 7 | 46, -14, 8 |
| Superior temporal gyrus (STG) | -61, -32, 8 | 59, -25, 8 |
| Primary somatosensory cortex (S1) | -53.4, -21.9, 60.7 | 47.9, -31.6, 63.9 |
| Secondary somatosensory cortex (S2) | -57.4, -27.7, 46 | 53.7, -38.1, 59.6 |
| Inferior frontal gyrus (IFG) | -46, 20, 8 | 46, 20, 8 |

Figure 1: **Source reconstruction.** Glass brain with MNI coordinates representing the sources included in the PEB analysis adjacent to a table with an overview of MNI coordinates and corresponding brain regions.

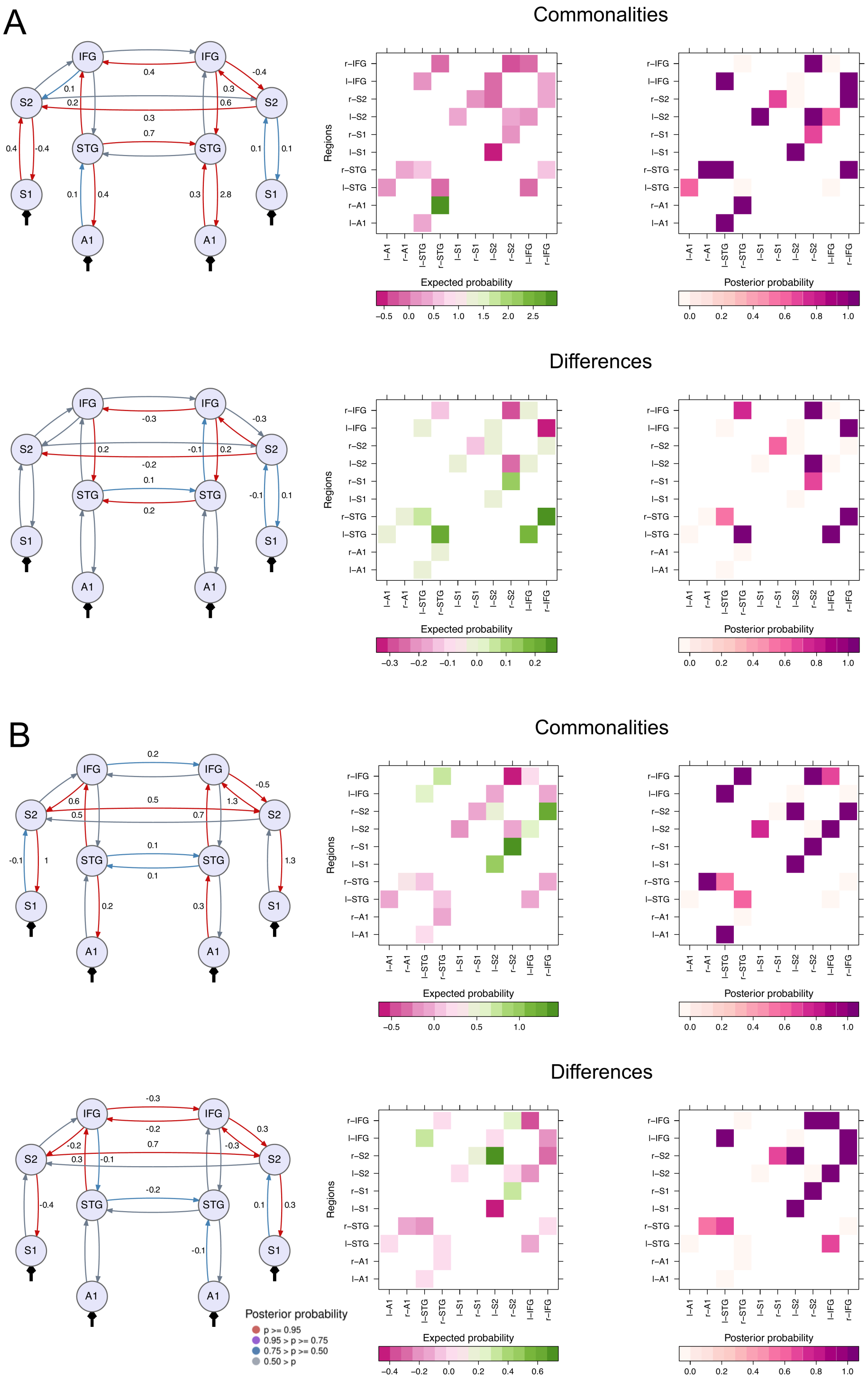


Figure 2: **Effective connectivity modelling.** We show commonalities and differences (B) between regular and salient deviants modelled using PEB for the auditory (A) and somatosensory condition (B). Arrows represent connections between cortical regions. Posterior expectations of connectivity parameters resulting from a multivariate normal probability density computation over PEB parameters that plotted next to each connection. Only extrinsic connections thresholded at a 95% posterior probability based on free energy (corresponding to very strong evidence) are shown. Below the glass brains, we show a heatmap with the posterior expected connectivity parameters on the left, and a heatmap with the corresponding posterior probabilities on the right.
